# Decreased miRNA-148a-3p expression in skeletal muscle of patients with chronic kidney disease

**DOI:** 10.1101/2022.05.24.493194

**Authors:** KA Robinson, LA Baker, MPM Graham-Brown, RU Ashford, Izabella Pawlyckz, RW Major, JO Burton, N. Sylvius, A Cooper, A Philp, EL Watson

## Abstract

**Introduction:** Skeletal muscle wasting is a common complication of chronic kidney disease which leads to a loss of muscle function. The pathogenesis of skeletal muscle wasting is incompletely understood, which is preventing the development of targeted therapeutics. Recent evidence implicates miRNAs in the of skeletal muscle wasting. Our aim was to firstly examine miRNA profiles of CKD human skeletal muscle for the identification of aberrant expression patterns compared to a healthy control (HC) cohort, and secondly, investigate the role these miRNAs may play in inducing or promoting skeletal muscle atrophy using a novel human primary skeletal muscle cell model of CKD skeletal muscle.

**Methods:** For the comparison between CKD and HC populations, skeletal muscle biopsies were collected from the vastus lateralis of n=15 non-dialysis dependent CKD patient’s stage 3b-5 CKD patients, and n=15 healthy controls matched for age, gender and physical activity. n=5 biopsies from each group underwent next generation sequencing to obtain complete microRNA profiles in CKD vs HC cohorts, which were then validated in a separate cohort by PCR (N=10 in each group). A causative role in muscle wasting was determined by transfection of key microRNAs into a primary culture model of CKD skeletal muscle and changes in protein degradation determined by L-[3H]- phenylalanine release into the media.

**Results:** Next Generation Sequencing identified differential expression of 16 miRNAs in skeletal muscle of CKD patients versus controls, and PCR validation confirmed miRNA-148a-3p expression was significantly decreased in CKD patients. The reduced miRNA-148a-3p expression was also maintained in the primary culture model. Upon overexpression of miRNA-148a-3p in CKD myotubes, protein degradation rates were decreased non-significantly (p=0.28) by 16.3% compared to un-transfected CKD cells.

**Conclusion:** CKD was associated with a significant reduction in miRNA-148a-3p expression in skeletal muscle compared to non-CKD controls which was retained in our *in vitro* model. Overexpression of miRNA-148a-3p in primary skeletal myotubes non-significantly decreased muscle protein degradation by 16.3%. In order to determine the importance of miRNA-148a-regulation of protein degradation, a deeper understanding of miRNA-148a-3p targets and their associated pathways with respect to those dysregulated in skeletal muscle wasting is required.

## Introduction

Skeletal muscle wasting is a common complication of chronic kidney disease (CKD), characterised by the rapid loss of muscle mass and strength, which leads to a loss of muscle function, physical inactivity, reduced quality of life^1^, and increased morbidity and mortality^2^. Notably, physical inactivity costs the UK approximately £7.4 billion per year, including costs of approximately £1.1 billion to the UK’s National Health Service (NHS) alone^3^. The association between skeletal muscle mass and mortality, and the high-cost burden to the NHS, makes this an important clinical consideration in the CKD population. However, despite extensive research, the pathogenesis of skeletal muscle wasting is incompletely understood, and therapeutic interventions are not in routine clinical practice. In order to develop therapeutic strategies with potential to improve these parameters, the key pathophysiological processes driving this complication need defining more completely.

A novel mechanism that may be involved in driving skeletal muscle atrophy is the aberrant expression of a family of short (18-25 nucleotides) non-coding RNA molecules, known as microRNAs (miRNAs). They are implicated in nearly every biological pathway, fine-tuning gene expression through binding to the 3’ untranslated region of target messenger RNAs (mRNAs) inducing either translational repression or degradation. Through this mechanism, several miRNAs can simultaneously regulate the expression of a single protein, whilst a single miRNA can target several different proteins, providing a layer of complexity to the regulation of gene expression^4^. Thus, miRNAs control the diversity of proteins within a cell, and consequently cellular activity.

Recent evidence implicates miRNAs in the development and progression of skeletal muscle wasting pathologies as crucial regulators of gene expression in skeletal muscle growth, regeneration, and metabolism^5^. Despite significant advancement in our understanding of the central role of miRNAs in regulating skeletal muscle wasting, a limited number of studies specifically investigate the role of miRNAs in CKD-associated muscle wasting, and current knowledge is restricted to animal models of CKD. These studies have reported the dysregulation of several miRNAs expressed within skeletal muscle, including miRNA-29, miRNA-486, miRNA-23a, miRNA-27a, and miRNA-26a^6-9^. Notably, therapeutic targeting of these miRNAs *in vivo* led to the prevention of CKD-induced skeletal muscle wasting and increased muscle mass despite the presence of CKD^6^. However, whilst miRNAs play a clear role in skeletal muscle wasting in animal models of CKD, there is a lack of evidence for their involvement in skeletal muscle wasting in human CKD. Knowledge of these pathways might enable the development of miRNA-based therapeutics for the treatment of muscle wasting in CKD and other catabolic conditions.

Our aim was to firstly examine miRNA profiles of CKD human skeletal muscle for the identification of aberrant expression patterns, and secondly, investigate the role these miRNAs may play in inducing or promoting skeletal muscle atrophy using a novel human primary skeletal muscle cell model of CKD skeletal muscle.

## Methods

### Study design and participants

Non-dialysis CKD patients stage G3-5 (eGFR <60mL/min/1.73m^2^) and non-CKD controls were recruited for a muscle biopsy as part of EXPLORE-CKD (ISRCTN18221837). Approval for this study was granted by the UK National Research Ethics Committee (reference 15/EM/0467), and all participants provided written informed consent. RNA-sequencing analysis was run on a discovery cohort of n=5 CKD patients and n=5 non-CKD controls. Results were subsequently validated in a separate validation cohort of n=10 CKD patients and n=10 non-CKD controls.

### Muscle biopsy procedure

Participants were matched for age, gender, and physical activity levels determined by the General Practice Physical Activity Questionnaire. Participants were excluded from undergoing a muscle biopsy if receiving warfarin or clopidogrel. Other exclusion criteria included: age <18years, pregnancy, insufficient command of English, or an inability to give informed consent. Patients were recruited from nephrology outpatient clinics at Leicester General Hospital, UK between September 2017 and February 2019. All patient biopsies were taken from the vastus lateralis (VL) using the microbiopsy technique^10^. Non-CKD control participants were recruited from orthopaedic theatre lists for the removal of benign intramuscular tumour using the open biopsy technique^11^. Samples were immediately dissected of any visible fat and connective tissue and placed into liquid nitrogen and stored until subsequent analysis.

### Small RNA-sequencing

CKD patient or non-CKD control biopsies (n=5) were used for small RNA-sequencing analysis. Briefly, total RNA was extracted from approximately 10mg/wet weight skeletal muscle tissue using TRIzol™ Reagent (Invitrogen™, 15596018). The amount of total RNA was quantified using the NanoDrop™ (Thermo Scientific™), and RNA quality was checked on a Bioanalyzer 2100 (Agilent, UK). Indexed small RNA libraries were prepared using the Illumina TruSeq Small RNA Library Preparation Kit (Illumina, UK) according to the manufacturer’s standard protocol. All libraries were pooled at equimolar concentrations, and PCR fragments of approximately 147-base pairs, corresponding to small RNA molecules with adapters and indexes, were selected. The quality of the resulting pooled library was checked on a Bioanalyzer High Sensitivity DNA chip (Agilent, UK) and sequenced by 36bp-single-end sequencing on an Illumina Miseq sequencer, using a 50-cycle MiSeq Reagent Kit v2 (Illumina, UK). The sequencing run was spiked-in with 1-% PhiX library (Illumina, UK) to control for the hardware and software performance of the instrument. The quality of the sequencing run was checked using the Miseq sequencing analysis viewer and Miseq Reporter v2.5.1, and Fastq sequencing files were generated by Miseq Reporter v2.5.1. The miRNAs and all other small RNA species were identified using the MiSeq Reporter small RNA workflow, which includes the trimming of the adapters, alignment of the sequencing reads against the reference genome using Bowtie v0.12.8, and the counting of the small RNA species. Raw read counts were normalised to the total number of mapped reads associated with each sample. Differential expression analysis of miRNAs was performed using the R package EdgeR, and significantly differentially expressed miRNAs were determined using a t-test with p≤0.05 as a cut-off.

### Quantitative RT-PCR

RNA was extracted from approximately 10mg/wet weight skeletal muscle tissue and primary cells using TRIzol™ Reagent (Invitrogen™, 15596018) and miRNAs were reverse transcribed to cDNA using the Applied Biosystems™ TaqMan™ Advanced miRNA cDNA Synthesis Kit (Applied Biosystems, UK, A28007). All PCR was carried out on Applied Biosystems™ Quantstudio™ 6 Flex Real-Time PCR system, and miRNA-specific primers, probes and internal controls were supplied as TaqMan™ Advanced miRNA assays (Applied Biosystems, UK, A25576): 477814_mir (hsa-miR-148a-3p), and 478578_mir (hsa-let-7f-5p) was used as an internal control, which was determined to remain stable over experimental conditions (data not shown). The mean Ct value was calculated from duplicate reactions, and the expression of the gene of interest was normalised to the housekeeper gene (ΔCt) and calculated as the difference between the Ct values of the two genes (2-ΔCt).

### In silico analysis of predicted microRNA targets

Target genes for miRNA-148a-3p were identified and compared using three online target prediction algorithms, mirDB^12^, TargetScan 7.2^13^, and DIANA-microT-CDS v5.0^14,15^. Putative targets for miRNA-148a-3p that were identified by all three of these algorithms were uploaded into BinGO^16^ (in CytoScape for full gene ontology (GO) enrichment analysis combining ‘biological process’, ‘molecular function’, and ‘cellular component’ categories. Cytoscape was used to create a visualisation network, as previously described^17^.

### Muscle cell isolation and differentiation

Primary skeletal muscle cells were isolated from skeletal muscle biopsies from CKD patients and non-CKD controls as described previously^18^. Briefly, muscle biopsies were minced into small fragments and enzymatically digested in two incubations for 15 minutes and then 5 minutes at 37°C with Collagenase Type IV (20mg), Bovine Serum Albumin (50mg) and Trypsin-EDTA with gentle agitation. The digest was strained through a 70μm nylon filter and centrifuged at 800 x g for 7 minutes. Cells were washed with Ham’s F-10 Nutrient Mix with 1% Penicillin-Streptomycin, 1% Amphotericin B, and 2% FBS and then pre-plated on uncoated 9cm^2^ petri-dishes in for 3 hours at 37°C under humidified 95% air and 5% CO_2_ to separate myogenic and non-myogenic cells. The cell suspension was transferred to a collagen I-coated 25cm^2^ flask and kept at 37°C under humidified 95% air and 5% CO2 until cells achieved approximately 40% confluence, and at this point, cells were passaged to increase stocks.

For experimentation, muscle cells were cultured in Ham’s F-10 Nutrient Mix with 1% Penicillin-Streptomycin, 1% Amphotericin B, and 2% FBS (growth medium) in collagen I-coated 12-well plates at a density of 5×10^4^ cells/well. Differentiation was induced by switching cells from growth medium to differentiation medium (Dulbecco’s Modified Eagle Medium with 1% Penicillin-Streptomycin and 2% Horse serum), which was replaced every other day. Following 7 days in differentiation medium, muscle cells had differentiated and fused into myotubes, which were lysed in TRIzol™ Reagent (Invitrogen™, 15596018) for qPCR analyses or used in down-stream experiments.

### MicroRNA transfection

Myotubes were transfected using either 5nM hsa-miR-148a-3p miRCURY LNA miRNA Mimic (QIAGEN, YM00472598-ADB) or negative control miRCURY LNA miRNA Mimic (QIAGEN, YM00479902-ADB) with HiPerfect Transfection Reagent (QIAGEN, 301705) for 8 hours. After 8 hours, cells were washed with HBSS and lysed in TRIzol™ Reagent for qPCR analysis or used in protein degradation experiments.

### Protein degradation

Myotubes were pre-labelled with L-[3H]-phenylalanine (2μCi/mL of differentiation medium; PerkinElmer, U.S, NET1122001MC) for 64 hours at 37°C under humidified 95% air and 5% CO_2_. Cells were washed with HBSS to remove non-incorporated material and transfected as described above. Following transfection, cells were incubated in differentiation medium supplemented with 2mM L-phenylalanine. Each experiment included proteasome inhibitor 10mM MG132 as a negative control. Aliquots of 300μL of culture medium were taken at 17-, 24-, and 48-hours post-transfection and stored at 4°C for the quantification of L-[3H]-phenylalanine. After 48 hours, the culture medium was aspirated, and cells were washed 3 times in ice-cold 0.9% w/v NaCl. To determine residual 3H in the cell protein, 500μL 0.5M NaOH was added to each well, cells were mechanically dislodged, transferred to a new tube and incubated at 70°C for 30 minutes. Following incubation, 50μL of the NaOH digest was transferred to 4mL Ecoscint A Scintillant (National Diagnostics, Hessle, UK). Media aliquots were deproteinised using 300μL 20% w/v trichloroacetic acid (TCA), incubated for 30 minutes at 4°C to precipitate proteins and subsequently centrifuged at 3,500rpm for 10 minutes at 4°C. Following centrifugation, 500μL of the TCA-soluble supernatant was added to 4mL Ecoscint A Scintillant and vortexed for the measurement of 3H using liquid scintillation counting. Total 3H is the sum of the residual radioactivity in cell proteins and the TCA-soluble 3H at the different time points. Protein degradation rates are expressed as log10 of the percentage of the total 3H remaining in cells per hour.

### Statistical analysis

All data in tables are represented as median (interquartile range). All data in figures are represented as mean ± standard error of the mean. Data were tested for normal distribution using the Shapiro Wilk test. Non-normally distributed data were either log-transformed prior to analysis or a non-parametric equivalent was used as appropriate. For un-paired group analysis, Independent Samples T-tests were performed for normally distributed data, and non-parametric Independent Samples Mann-Whitney U tests were performed for non-normally distributed data. For paired comparisons, Paired Samples T-tests were performed for normally distributed data, and non-parametric Wilcoxon tests were performed for non-normally distributed data. For comparisons involving three or more groups, a one-way ANOVA with Tukey’s multiple comparison test was performed for normally distributed data, and a Games-Howell test was performed for non-normally distributed data. Statistical significance was accepted at p≤0.05. All statistical analysis was carried out using IBM SPSS Statistics Version 26 (IBM, Chicago, IL).

## Results

### Participant characteristics

Skeletal muscle biopsies were collected from a total of 15 CKD patients and 15 matched non-CKD controls. Skeletal muscle biopsies from 5 CKD patients and 5 matched healthy controls were included in the small RNA-sequencing analysis, and the individual characteristics of this small RNA-sequencing cohort can be found in Table 1. Skeletal muscle biopsies from 10 CKD patients and 10 matched healthy controls were used in the qPCR validation of miRNA expression, and the individual characteristics of this validation cohort can be found in Table 2. Skeletal muscle biopsies from a sub-group of the validation cohort (n = 7 per group) were used in primary cell experiments for the functional analysis of miRNAs *in vitro*, and the individual characteristics of this primary cell cohort can be found in Table 3.

**Table 1.**
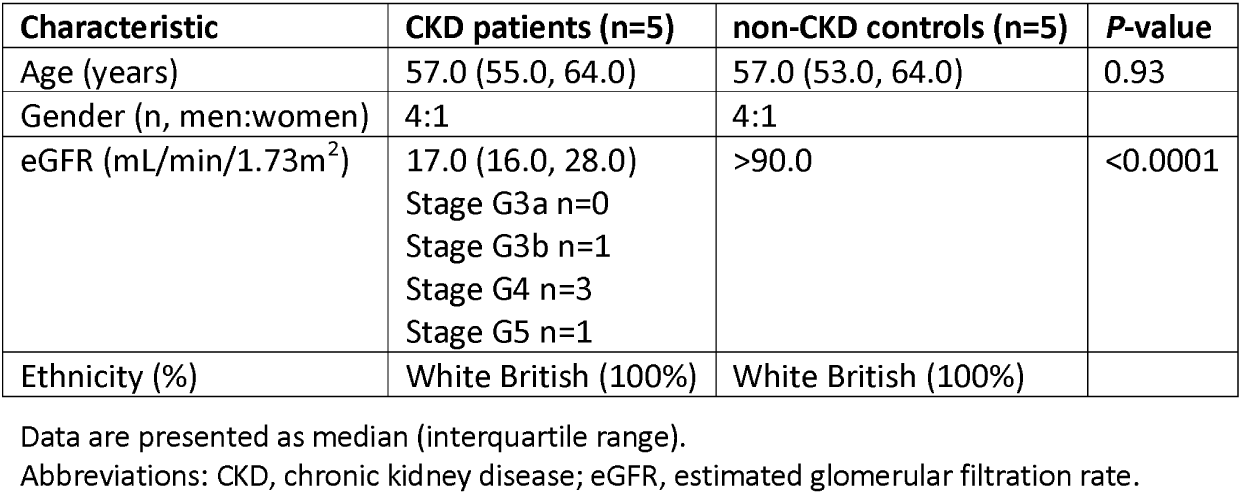
Characteristics of CKD patients and non-CKD controls in small RNA-sequencing cohort

| Characteristic | CKD patients (n=5) | non-CKD controls (n=5) | P-value |
| --- | --- | --- | --- |
| Age (years) | 57.0 (55.0, 64.0) | 57.0 (53.0, 64.0) | 0.93 |
| Gender (n, men:women) | 4:1 | 4:1 |  |
| eGFR (mL/min/1.73m<sup>2</sup>) | 17.0 (16.0, 28.0)<br>Stage G3a n=0<br>Stage G3b n=1<br>Stage G4 n=3<br>Stage G5 n=1 | >90.0 | <0.0001 |
| Ethnicity (%) | White British (100%) | White British (100%) |  |

Data are presented as median (interquartile range).
Abbreviations: CKD, chronic kidney disease; eGFR, estimated glomerular filtration rate.

**Table 2.** Characteristics of CKD patients and non-CKD controls in validation cohort

| Characteristic | CKD patients (n=10) | Non-CKD controls (n=10) | P-value |
| --- | --- | --- | --- |
| Age (years) | 62.0 (55.2, 70.5) | 66.0 (55.0, 71.5) | 0.99 |
| Gender (n men/women) | 8:2 | 8:2 |  |
| eGFR (mL/min/1.73m<sup>2</sup>) | 28.0 (20.2, 41.0)<br>Stage G3a n=2<br>Stage G3b n=3<br>Stage G4 n=4<br>Stage G5 n=1 | 82.5 (80.2, 85.5) | <0.0001 |
| Physically inactive (%) | 55% | 50% | 0.61 |

Data are presented as median (interquartile range).
Abbreviations: CKD, chronic kidney disease; eGFR, estimated glomerular filtration rate.
Physical activity status was determined using the GPAQ Questionnaire.

**Table 3.** Characteristics of CKD patients and non-CKD controls in primary cell cohort

| Characteristic | CKD patients (n=7) | non-CKD controls (n=7) | P-value |
| --- | --- | --- | --- |
| Age (years) | 57.0 (47.5, 68.5) | 45.0 (39.5, 66.5) | 0.53 |
| Gender (n men/women) | 3:4 | 3:4 |  |
| eGFR (mL/min/1.73m<sup>2</sup>) | 22.0 (15.5, 32.5)<br>Stage G3a n=0<br>Stage G3b n=2<br>Stage G4 n=3<br>Stage G5 n=2 | 81.0 (74.0, 90.0) | <0.0001 |

Data are presented as median (interquartile range).
Abbreviations: CKD, chronic kidney disease; eGFR, estimated glomerular filtration rate.

### CKD decreases miRNA-148a-3p expression in skeletal muscle

Small RNA-sequencing identified differential expression of 16 miRNAs (fold-change values of ≥1.5) in CKD muscle compared to non-CKD controls (Table 4). Upon validation of miRNAs by qPCR, we identified a significant 1.3-fold decrease was maintained in the level of miRNA-148a-3p (p=0.03) in the skeletal muscle of patients with CKD versus non-CKD controls (Fig 1a). No further significant differences were maintained upon validation (data not shown). To determine whether decreased miRNA-148a-3p is skeletal muscle specific, we measured miRNA-148a-3p in primary skeletal muscle cells isolated from patients with CKD and healthy controls during proliferation (0 days; 0D), early differentiation (3 days; 3D) and late differentiation (7 days; 7D; Fig 1b). In these cells, miRNA-148a-3p was significantly decreased threefold (p=0.04) in CKD myotubes (7D) compared to those from non-CKD controls (Fig 1b). miRNA-148a-3p expression was also decreased in CKD derived myoblasts at 0D and 3D, however this did not reach statistical significance (Fig 1b). All later experiments were therefore carried out in mature skeletal muscle myotubes.

**Table 4.** Sixteen skeletal muscle miRNAs with differential expression in CKD patients compared to non-CKD controls as determined by small RNA-sequencing

| miRNA | Fold-change (Increased) | Fold-change (Decreased) |
| --- | --- | --- |
| miRNA-126-3p | 1.98 |  |
| miRNA-128-3p | 1.54 |  |
| miRNA-148a-3p | 2.09 |  |
| miRNA-182-5p | 1.45 |  |
| miRNA-21-5p | 1.80 |  |
| miRNA-22-3p | 1.59 |  |
| miRNA-29c-3p | 1.52 |  |
| miRNA-92a-3p | 1.87 |  |
| miRNA-100-5p |  | 2.22 |
| miRNA-191-5p |  | 1.55 |
| miRNA-206 |  | 1.62 |
| miRNA-486-5p |  | 1.60 |
| miRNA-99a-5p |  | 1.57 |
| miRNA-99b-5p |  | 1.93 |
| Let-7a-5p |  | 1.48 |
| Let-7e-5p |  | 1.54 |

**Fig 1.**
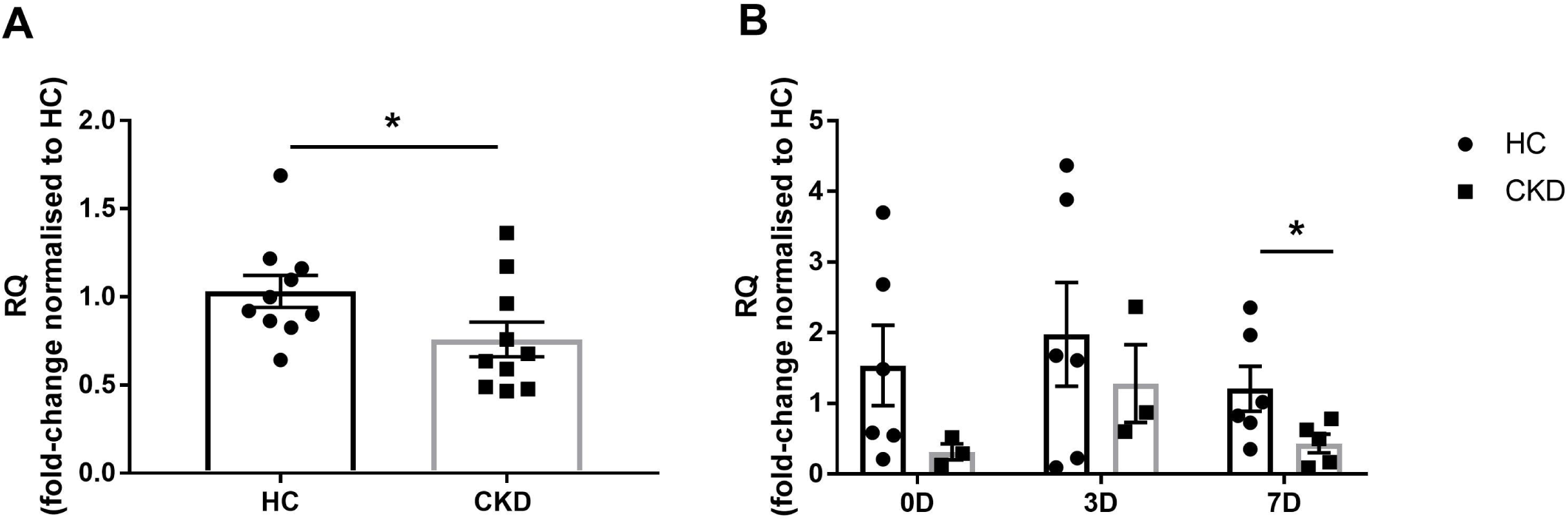
CKD results in decreased miRNA-148a-3p expression within skeletal muscle. Data are presented as mean ± SEM. Real time PCR data expressed using the RQ method relative to matched healthy controls (HC); results are normalised to Let-7f as an internal control. (A) miRNA-148a-3p expression in lower limb skeletal muscle biopsies of CKD patients and age-and sex-matched healthy controls (HC). * Denotes *P* = 0.03 vs healthy controls. (B) miRNA-148a-3p expression in human primary skeletal muscle cells from CKD patients and matched healthy controls. Expression was determined at three timepoints representing proliferation (0D; 0 days), early differentiation (3D; 3 days) and late differentiation (7D; 7 days). * Denotes *P* = 0.048 vs healthy controls.

### *In silico* analysis of miRNA-148a-3p predicted targets

To gain an insight into the functional role of miRNA-148a-3p, miRNA-148a-3p targets were investigated using online target prediction databases, miRDB, TargetScan, and DIANA-microT-CDS. There were 361 genes predicted by all three databases, which were uploaded into BinGO in CytoScape for full GO enrichment analysis (Table 5). GO analysis indicated that miRNA-148a-3p is involved in the regulation of cellular metabolism, and as skeletal muscle wasting is a condition of disordered muscle metabolism, this highlights a promising role for miRNA-148a-3p in skeletal muscle wasting.

**Table 5.** Gene ontology (GO) enrichment analysis of miRNA-148a predicted targets

| GO Description | P-value |
| --- | --- |
| regulation of biological process | 2.50E-16 |
| biological regulation | 4.38E-16 |
| regulation of cellular process | 8.79E-16 |
| protein binding | 3.51E-15 |
| binding | 7.31E-15 |
| positive regulation of cellular process | 4.62E-12 |
| positive regulation of biological process | 6.48E-12 |
| cellular macromolecule metabolic process | 6.30E-11 |
| regulation of metabolic process | 8.46E-11 |
| regulation of cellular metabolic process | 7.65E-10 |

### Experimental validation of miRNA-148a-3p predicted targets

Due to the central role of disordered muscle metabolism in the pathogenesis of CKD skeletal muscle wasting, miRNA-148a-3p predicted targets from the GO category cellular macromolecule metabolic process, in addition to targets with proven roles in the maintenance of muscle mass, and in muscle atrophy, were selected for experimental validation. There were no significant differences in expression of miRNA-148a-3p targets in skeletal muscle in CKD patients compared to healthy controls (Figure4a). However, small increases in LDLR (1.34-fold; p=0.079), MEOX2 (1.10-fold; p=0.387), PNRC2 (1.15-fold; p=0.315) and USP6 (1.19-fold; p=0.579) were observed (Figure4a).

The expression of these genes was examined *in vitro* in human primary skeletal myotubes with and without miRNA-148a-3p overexpression. No differences in expression were observed for LDLR (p=0.98), MEOX2 (p=1.00) and PNRC2 (p=0.91) in CKD myotubes compared to controls (Figure4b). The expression of USP6 was undetermined and therefore, was not included in the analysis. Moreover, miRNA-148a-3p overexpression did not alter gene expression in CKD myotubes (LDLR p=0.50; MEOX2 p=0.94; PNRC2 p=0.68; Figure4b).

### miRNA-148a-3p overexpression and protein degradation in CKD myotubes

Evidence strongly suggests that the persistent activation of muscle protein degradation via the ubiquitin proteasome system (UPS) is the primary cause of muscle wasting in CKD^19^. To assess if there is a role of miRNA-148a-3p in the regulation of muscle protein degradation, we transfected CKD myotubes with a miRNA-148a-3p mimic or control mimic. miRNA-148a-3p expression was significantly increased in CKD myotubes transfected with miRNA-148a-3p mimic compared to cells transfected with control mimic at the same concentration (p=0.02; Fig 2).

**Fig 2.**
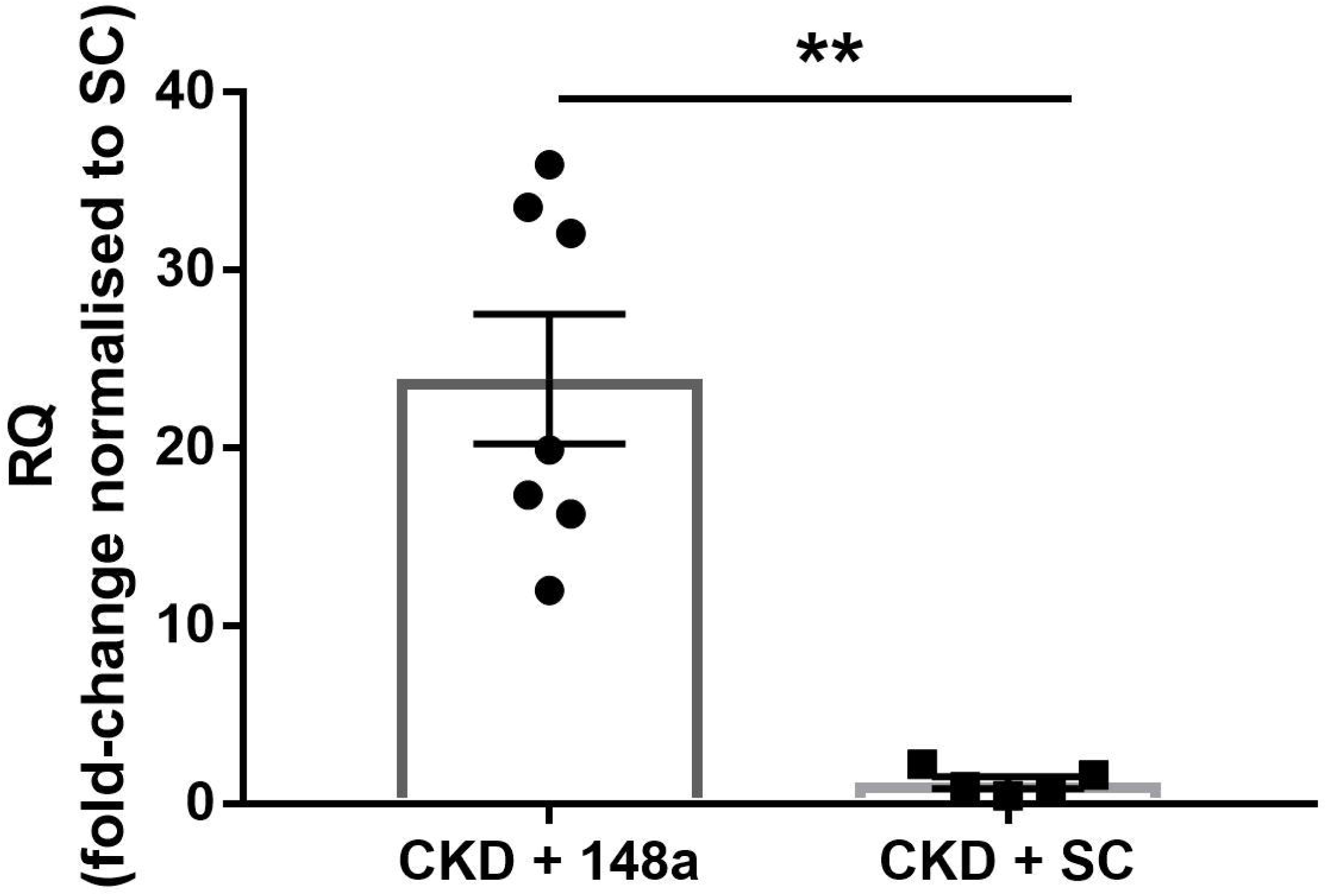
miRNA-148a-3p expression in human primary skeletal myotubes from CKD patients after the addition of miRNA-148a-3p mimic (148a) or control mimic (SC) at 5nM concentrations for 8 hours to CKD myotubes. Data are presented as mean ± SEM. Real time PCR data expressed using the RQ method relative to control mimic; results are normalised to Let-7f as an internal control. ** Denotes *P* = 0.02 vs control mimic.

Basal rates of protein degradation were significantly higher in CKD derived myotubes (1.3-fold or 29.3% increase) compared to non-CKD controls (p=0.032; Fig 3). Because decreased miRNA-148a-3p expression and increased protein degradation are maintained in CKD primary skeletal muscle cells *in vitro*, we hypothesised that enhancing miRNA-148a-3p expression would decrease protein degradation rates. Upon overexpression of miRNA-148a-3p in CKD myotubes, protein degradation rates were decreased by 16.3% compared to un-transfected CKD cells, however this reduction did not reach statistical significance (p=0.28; Fig 3). Following transfection, the protein degradation rates were now only 7.5% higher than the non-CKD controls, with no statistical difference between the groups (Fig 3). Interestingly, the control mimic also decreased protein degradation rates by 9.4% compared to un-transfected CKD cells (p=0.75; Fig 3). However, this decrease was not significant and not to the same extent as that seen with miRNA-148a-3p mimic. As expected, proteasome inhibitor, MG132, significantly decreased protein degradation rates compared to all conditions (p≤0.001; Fig 3). These results suggest that miRNA-148a-3p overexpression reduced protein degradation rates in CKD myotubes, albeit non-significantly.

**Fig 3.**
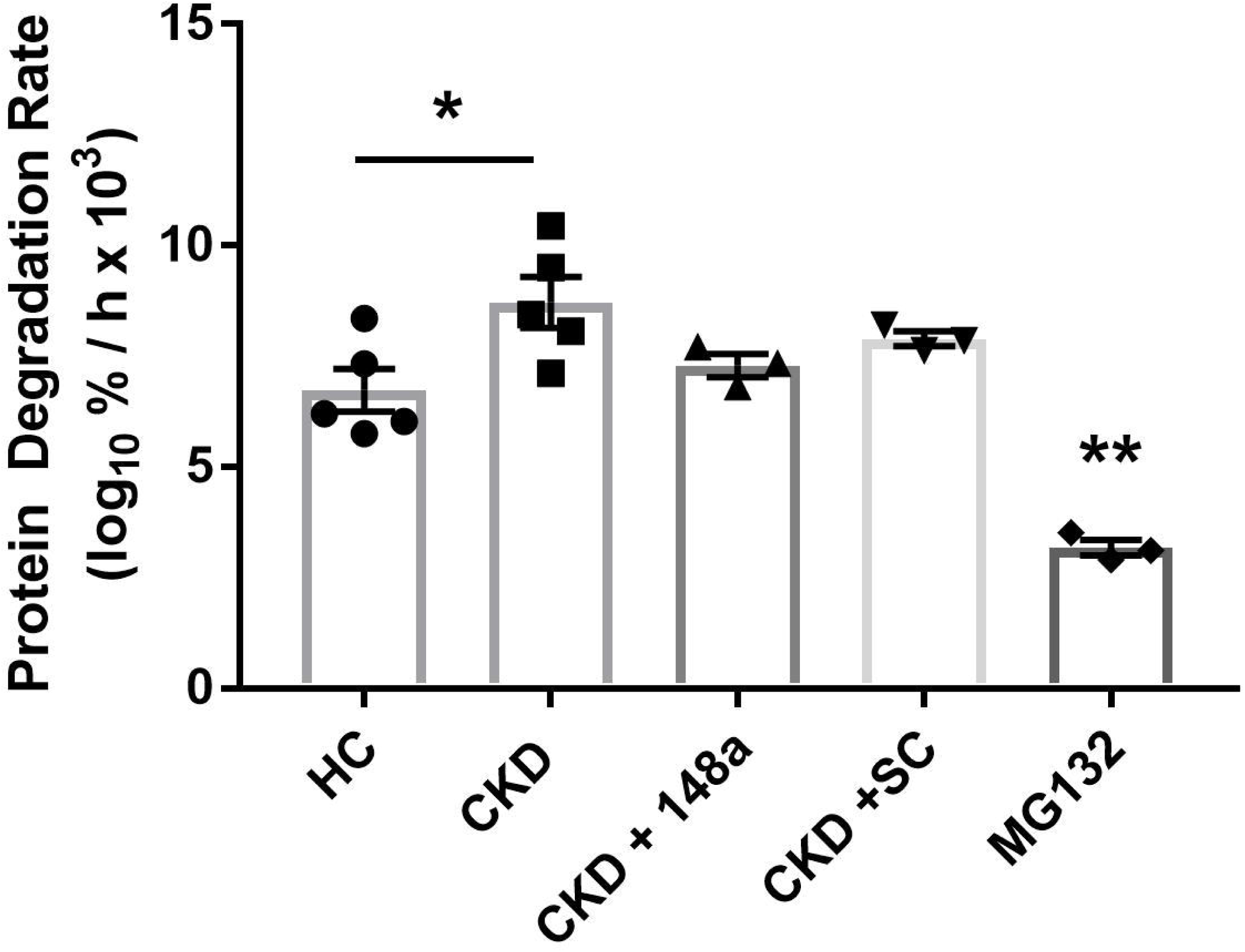
Protein degradation rates in human primary skeletal myotubes from CKD patients and healthy controls (HC). Data are presented as mean ± SEM. All rates were determined by linear regression over a time course of 17-48 h after the addition of miRNA-148a-3p mimic (148a) or control mimic (SC) at 5nM concentrations for 8 hours. A proteasome inhibitor (MG132) was included as a negative control. * Denotes *P* = 0.032 vs healthy controls; ** Denotes *P* ≤0.001 vs all conditions.

**Fig 4.**
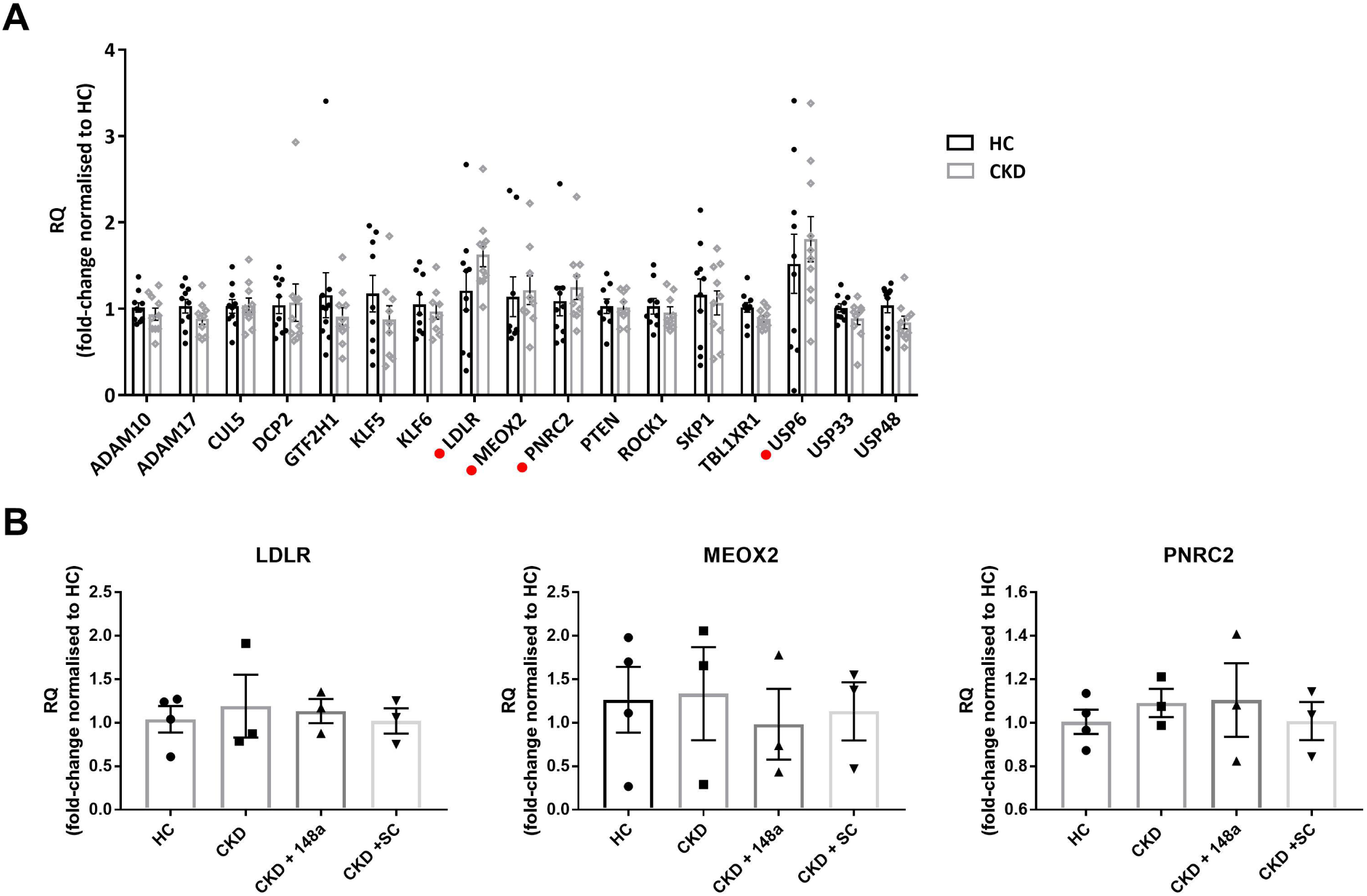
Expression of miRNA-148a-3p predicted targets measured by real-time quantitative PCR. (A) skeletal muscle biopsies from CKD patients and matched healthy controls (HC). The bar graph shows gene expression calculated using the RQ method, results are normalised to RPLP0 as an internal control (bars mean ± SEM; n=10) C indicates genes of interest. (B) Expression of miRNA-148a-3p predicted targets in human primary skeletal myotubes from CKD patients and matched HCs. Gene expression was measured after the addition of miRNA-148a-3p mimic (148a) or control mimic (SC) at 5nM concentrations for 8 hours. The bar graph shows gene expression calculated using the RQ method, results are normalised to RPLP0 as an internal control (bars mean ± SEM; HC n=4; CKD n=3; CKD + 148a n=3; CKD + SC n=3).

## Discussion

Here we provide the first evidence that skeletal muscle miRNA expression is dysregulated in the skeletal muscle of human patients with non-dialysis CKD, and that aberrant miRNA expression may contribute in part to the increased protein degradation rate that is closely linked to a loss of muscle mass. We report that CKD was associated with a significant reduction in miRNA-148a-3p expression in skeletal muscle compared to non-CKD controls and that miRNA-148a-3p overexpression in a novel primary cell model of human skeletal muscle cells isolated from CKD patients and healthy controls^18^ non-significantly decreased muscle protein degradation by 16.3%. Subsequent miRNA-148a-3p target analysis revealed that miRNA-148a-3p targets are enriched in processes of cellular metabolism, highlighting a potential undiscovered role for miRNA-148a-3p in the modulation of abnormal cellular metabolism, which drives the development and progression of skeletal muscle wasting in CKD.

The miRNA-148/152 family consists of three members: miR-148a, miR-148b and miR-152, which are located on chromosomes 7, 12 and 17, respectively^20^. The role of the miR-148/152 family has been described in several biological processes, and its aberrant expression has been frequently observed in many pathologies including IgA nephropathy^21^, lupus nephritis^22^, type 1 diabetes^23^, atherosclerosis^24^, chronic fatigue syndrome^25^ and several cancers^26^. Specifically, miRNA-148a has previously been identified as a novel myogenic miRNA that is highly expressed in skeletal muscle and of importance in the regulation of myogenesis in mouse^27^, chicken^28^, and bovine^29^ models. For example, increased miRNA-148a expression promoted myoblast differentiation in C2C12 myoblasts, mouse primary skeletal myoblasts and chicken skeletal muscle. In these models, miRNA-148a inhibited Rho-associated protein kinase 1 (ROCK1), a known inhibitor of myogenesis, and of Mesenchyme homeobox 2 (MEOX2), respectively. Yin et al. further demonstrated that the effects of miRNA-148a on myogenic differentiation was mediated via increased p-Akt levels and subsequent activation of the PI3K-Akt signalling pathway, which is critical in the maintenance of skeletal muscle mass and is disturbed in CKD^30^. Increased miRNA-148a has also been shown to regulate PI3K-Akt signalling in the setting of atherosclerosis via the suppression of FoxO3 mRNA expression in endothelial cells^24^. Thus, if the same holds true in human skeletal muscle, the ability of increased miRNA-148a to encourage activation of the PI3K-Akt signalling pathway through both increased p-Akt levels, as well as decreased FoxO3 mRNA expression, could have promising therapeutic potential in CKD-associated skeletal muscle wasting.

Other pathways regulated by miRNA-148a that are relevant to skeletal muscle wasting include the TGF-β pathway and the autophagy-lysosomal pathway, which might provide a potential link between miRNA-148a dysregulation and skeletal muscle wasting. There is an abundance of evidence that TGF- β and members of the TGF-β family (such as myostatin) are closely associated with muscle-protein loss in catabolic conditions via activation of Smad2/Smad3-mediated signalling pathway in skeletal muscle, which leads to stimulation of proteolysis and muscle atrophy^31^. There are several studies that confirm a role for miRNA-148a in the regulation of TGF-β/Smad signalling. For example, it was reported that overexpression of miRNA-148a promoted a miRNA-148a-mediated inhibition of the TGF-β/Smad2 signalling pathway in gastric cancer cells^32^ and in hepatocellular carcinoma both *in vitro* and *in vivo*^33,34^. In contrast however, Wang et al. reported that miRNA-148a expression in glioblastoma led to an enhanced-strength and prolonged-duration of TGF-β/Smad activation in mice, suggesting that miRNA-148a might have different effects in differing cell types^35^. These findings were also consistent with a significant correlation between miRNA-148a levels and activated TGF-β/Smad signalling in a cohort of human glioblastoma specimens. Overall, these studies highlight a role for miRNA-148a in the regulation of TGF-β signalling, a key pathway in regulating skeletal muscle wasting. Indeed, downregulation of skeletal muscle miRNA-148a expression in CKD patients could be perpetuating the activation of TGF-β/Smad2 signalling and contributing to skeletal muscle atrophy. Therefore, it is possible that miRNA-148a overexpression might inhibit TGF-β/Smad2 signalling to potentially reduce skeletal muscle atrophy in this population.

It has been proven that miRNA-148a can regulate key pathways that stimulate protein degradation including the PI3K-Akt, TGF-β, and autophagy/lysosome pathways, however, the role of miRNA-148a has not yet been explored with respect to skeletal muscle atrophy in CKD or other wasting conditions. Evidence strongly suggests that persistent activation of protein degradation is the primary cause of muscle protein breakdown in CKD, and in the present study, protein degradation rates were significantly increased by 29.3% in CKD myotubes compared to non-CKD controls. Overexpression of miRNA-148a-3p reduced protein degradation rates in CKD myotubes by 16.3%, however, this did not reach statistical significance. Indeed, it could be reasoned that a 16.3% reduction in protein degradation rates has the potential to reduce the severity of muscle atrophy in patients with CKD as evidence suggests that even small increases in protein degradation over time result in substantial skeletal muscle wasting^30^.

Alternatively, this could suggest that miRNA-148a-3p either does not adequately suppress all of the key factors driving muscle proteolysis, such as insulin resistance, myostatin, and inflammation or does not impact pathways central to the regulation of protein degradation, but still contributes to an extent. In the present study, miRNA-148a targets were not identified in skeletal muscle, and further exploration of targets is required to gain an understanding of the mechanism through which miRNA-148a-3p regulates protein degradation.

In conclusion, CKD was associated with a significant reduction in miRNA-148a-3p expression in skeletal muscle compared to non-CKD controls which was retained in our *in vitro* model. Overexpression of miRNA-148a-3p in primary skeletal myotubes non-significantly decreased muscle protein degradation by 16.3%. In order to determine the importance of miRNA-148a-regulation of protein degradation, a deeper understanding of miRNA-148a-3p targets and their associated pathways with respect to those dysregulated in skeletal muscle wasting is required. However, evidence for miRNA-148a regulation of pathways that are central to skeletal muscle wasting, including PI3K-Akt and TGF-β pathways, suggests a key role for miRNA-148a in skeletal muscle wasting in patients with CKD.

